# From the cage to the wild: Introductions of Psittaciformes to Puerto Rico with emphasis on the invasive ecology of the white-winged parakeet

**DOI:** 10.1101/264937

**Authors:** Wilfredo Falcón L., Raymond L. Tremblay

## Abstract

In this study, we assessed invasions of Psittaciformes in Puerto Rico. We reviewed the literature, public databases, citizen science records, and performed in situ population surveys across the island to determine the historical and current status and distribution of psittacine species. We used count data from *Ebird* to determine population trends. For species whose populations were increasing, we modelled their potential distribution using niche modeling techniques. Focusing on the white-winged parakeet *(Brotogeris versicolurus)*, which was considered the most successful psittacine species by the year 2000, we evaluated the population size, calculated growth rates and estimated the breeding proportion in two populations by performing roost counts for four consecutive years. We found a total of 46 Psittaciformes present in Puerto Rico, of which 26% are only present as pets, at least 29 species have been reported in the wild, and of those, there is evidence that at least 12 species are breeding. Our results indicate that most introduced species which have been detected as established still persist, although mostly in localized areas and small populations. Clear evidence of invasiveness was found for *B. versicolurus* and *Myiopsitta monachus*, which have greatly expanded their range. Moreover, *Psittacara erythrogenys* and *Eupsittacula canicularis* also showed population increase. The niche models predicted suitable areas for the four species, and also indicate the potential for range expansion. Population estimates of the white-winged parakeet during the study period showed a steady increase, and exhibited exponential growth, with geometric mean population growth rates of 1.25 per year. Currently growth rate of the white-winged parakeet does not appear to be limited by any predator, resources or nest availability, and we expect them to continue increasing and expanding their range. We discuss the factors leading to invasion success, assess the potential impacts, and we discuss possible management strategies and research prospects.

## Introduction

Over the last decades, the rate of species introductions has risen considerably, mirroring the rise of international trade, and the shifts in species distributions and the reorganization of biodiversity are now considered a signature of the Anthropocene (Seebens et al., 2017). In parallel, the study of invasive species has gained broad attention from ecologists, government agencies and the public due to the potential negative economical, ecological and environmental impact that these may cause on ecological processes and patterns (Sanders et al., 2003; Mooney, 2005; Davis, 2009; Lockwood, Hoopes & Marchetti, 2013). One of the factors that most contributed to the establishment of nonnative species in recent history has been the pet trade (Smith et al., 2009).

Among the species commonly sold as exotics are Psittaciformes (parrots, conures and cockatoos; hereon referred to collectively as parrots), with around two-thirds of species being transported outside their native range (Cassey et al., 2004b). It has been estimated that around USD $1.4 billion were generated during the 1990’s as a result of the global trade with Psittaciformes, and that around four million birds were captured from the wild to supply the demand of the pet trade industry (Thomsen et al., 1992). Four general modes of introductions have been identified for parrots: releases by traders due to oversupply or legal complications (Forshaw, 1973; Robinson, 2001), or the accidental or intentional releases by pet owners (Blackburn, Lockwood & Cassey, 2009).

Two thirds of successful avian introductions have been on islands (Blackburn, Lockwood & Cassey, 2009). Many species of Psittaciformes have been imported to be sold as pets in Puerto Rico, especially since the 1950’s and as many as eight species of parrots have been recorded as established (or probably established) by early 2000 (Pérez-Rivera, 1985; Raffaele, 1989; Camacho-Rodríguez, Chabert-Llompart & López-Flores, 1999; Oberle, 2000). In Puerto Rico, there is evidence that three endemic species of parrots have been present on the island, but the only extant species is the endemic Puerto Rican amazon *(Amazona vittata vitta)* and the two others species, *Amazona v. gracipiles* and *Psittacara maugei* having gone extinct (Snyder, Wieley & Kepler, 2007; Olson, 2015). As with any exotic and invasive species, local federal and state agencies are concerned with the possible effects that these species may have on indigenous species and the ecosystem, and managing agencies want to have information about the size, distribution and spread of exotic populations.

For example, the white-winged parakeet *(Brotogeris versicolurus)* has been considered the most successful psittacine species established in Puerto Rico (Pérez-Rivera et al., 1985; Raffaele, 1989; Camacho-Rodríguez, Chabert-Llompart & López-Flores, 1999; Oberle, 2000). Although Pérez-Rivera et al. (1985) alerted that the white-winged parakeet could expand and cause significant damage to crops, there is no evidence that this has yet materialized. Previous surveys has confirmed the persistence of locally common and self-sustaining populations of the white winged parakeet (Pérez-Rivera et al., 1985; Raffaele, 1989; Camacho-Rodríguez, Chabert-Llompart & López-Flores, 1999; Oberle, 2000). Since 2004, and 35 years after it was first reported in the wild in Puerto Rico, the population in the Metropolitan Area of San Juan appeared to be expanding aggressively. This brought renewed concerns about the possible negative effects that the parakeet might have on the native flora and fauna if the white-winged parakeet kept expanding its range, and on the agricultural sector.

In this study, we present a review on the introduction and persistence of Psittaciformes in Puerto Rico, and evaluated their historic and present distribution. In more detail, we assess the invasive ecology of the white-winged parakeet – including aspects of their natural history, population estimates, and population growth rates. Also, we present the predicted distribution of white-winged parakeets on the island based on niche modelling techniques. Finally, we address possible factors that may have contributed to the establishment success of species of Psittaciformes in Puerto Rico, and discuss our results in the context of management and prospects.

## Materials and Methods

### Introductions of Psittaciformes to Puerto Rico

Members of the Psittaciformes, which comprise about 393 species in 92 genera that include macaws, cockatoos, parrots, and conures, are mostly pantropical, although some species inhabit temperate areas in the southern hemisphere (Juniper & Parr, 1998; Forshaw, 2010). They are considered one of the most endangered groups of birds in the world, and threats include trapping for trade, habitat destruction and hunting (Snyder et al., 2000).

#### Historical and current status of Psittaciformes

To assess the historical introductions of Psittaciformes to Puerto Rico, and their current status, we surveyed historical reports on the distribution of the species (Forshaw, 1973;Pérez-Rivera & Vélez-Miranda, 1980; Pérez-Rivera, 1985; Pérez-Rivera et al., 1985; Raffaele, 1989; Camacho-Rodríguez, Chabert-Llompart & López-Flores, 1999; Oberle, 2000). We also recorded species based on observations performed during 2013–2017. In addition, we surveyed the *E-bird* (http://ebird.com/) online database, which contain records from citizen-science-derived reports and from experts. Moreover, performed searches on online local birding groups for photographic records using the search terms ‘parrot’, ‘parakeet’, ‘macaw’, ‘cockatoo’ in English and Spanish. The local groups included: *Aves de Puerto Rico* (https://www.facebook.com/avesdepuertoricoFelPe/), the *Puerto Rico Ornithological Society* (https://www.facebook.com/sociedadornitologicapuertorriquena/), *Bird Photographers of Puerto Rico* (https://www.facebook.com/groups/615958701756859/), and *Biodiversidad de Puerto Rico* (https://www.facebook.com/groups/PRNatural/). This was done between November–December 2017 (see electronic supplementary information S1 for the origin of the data). To identify species sold as pets but not seen in the wild, we visited pet stores, mainly in the Metropolitan Area of San Juan, and the pets section of local online classified ads *Clasificados Online* (www.clasificadosonline.com). Moreover, the online databases *Ebird, CABI Invasive Species Compendium* (https://www.cabi.org/isc) and the *Global Invasive Species Database* (http://www.iucngisd.org/gisd) were used to assess the invasiveness of species of Psittaciformes have reached in their non-native range (outside Puerto Rico) in the categorization scheme of invasive species according to (Blackburn et al., 2011; see Table 1). We also used the latter scheme to classify the status (invasiveness) of Psittaciformes in Puerto Rico. We used Forshaw (2010) for taxonomical classification and common names of Psittaciformes, the native distribution, and include any classification changes according to del Hoyo et al. (2014). The IUCN Red List (ver. 3.1; www.iucnredlist.org) was used to assess the conservation status for each species, population trends in their native range, and possible threats or reasons for population increase.

**Table 1:**
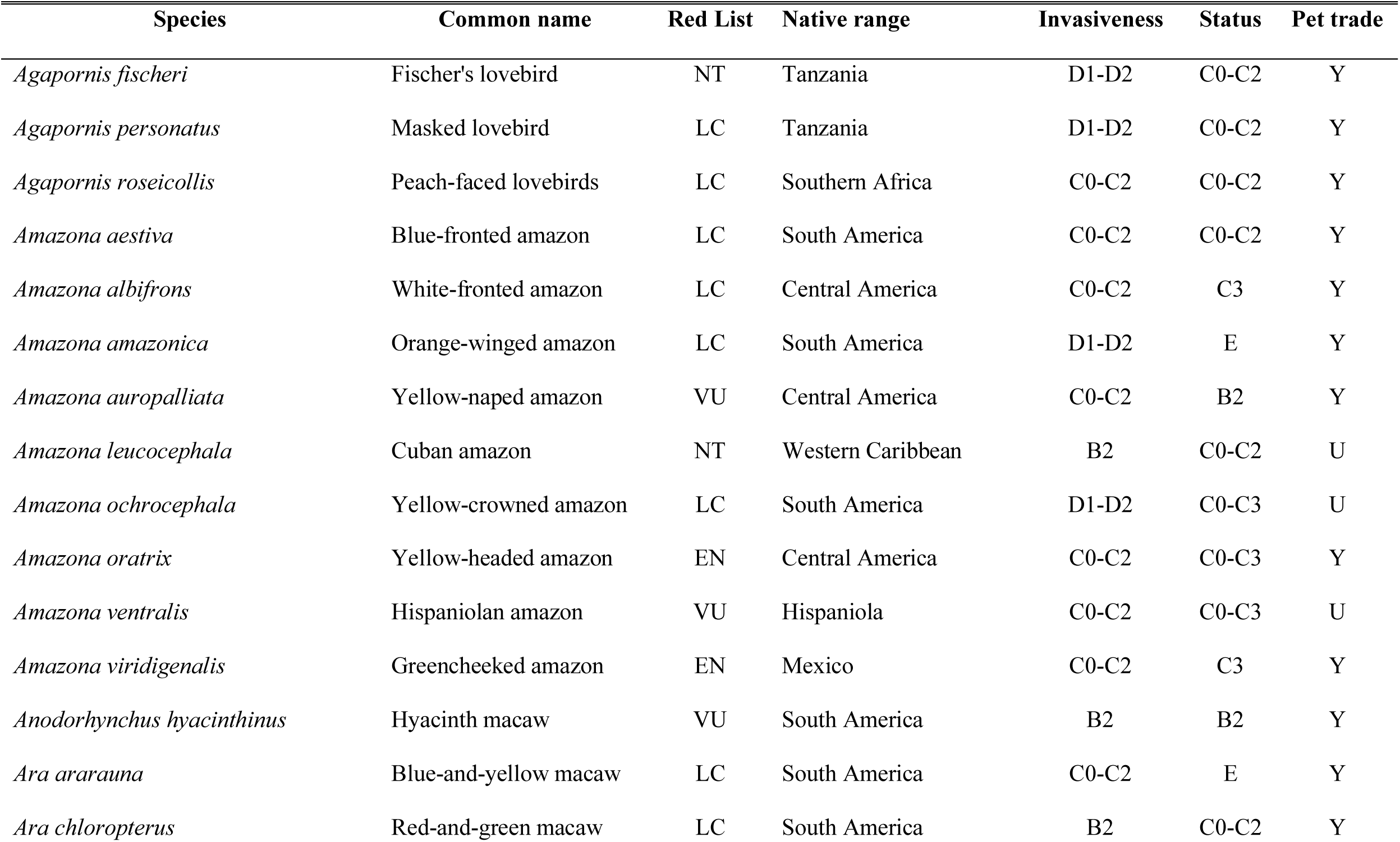
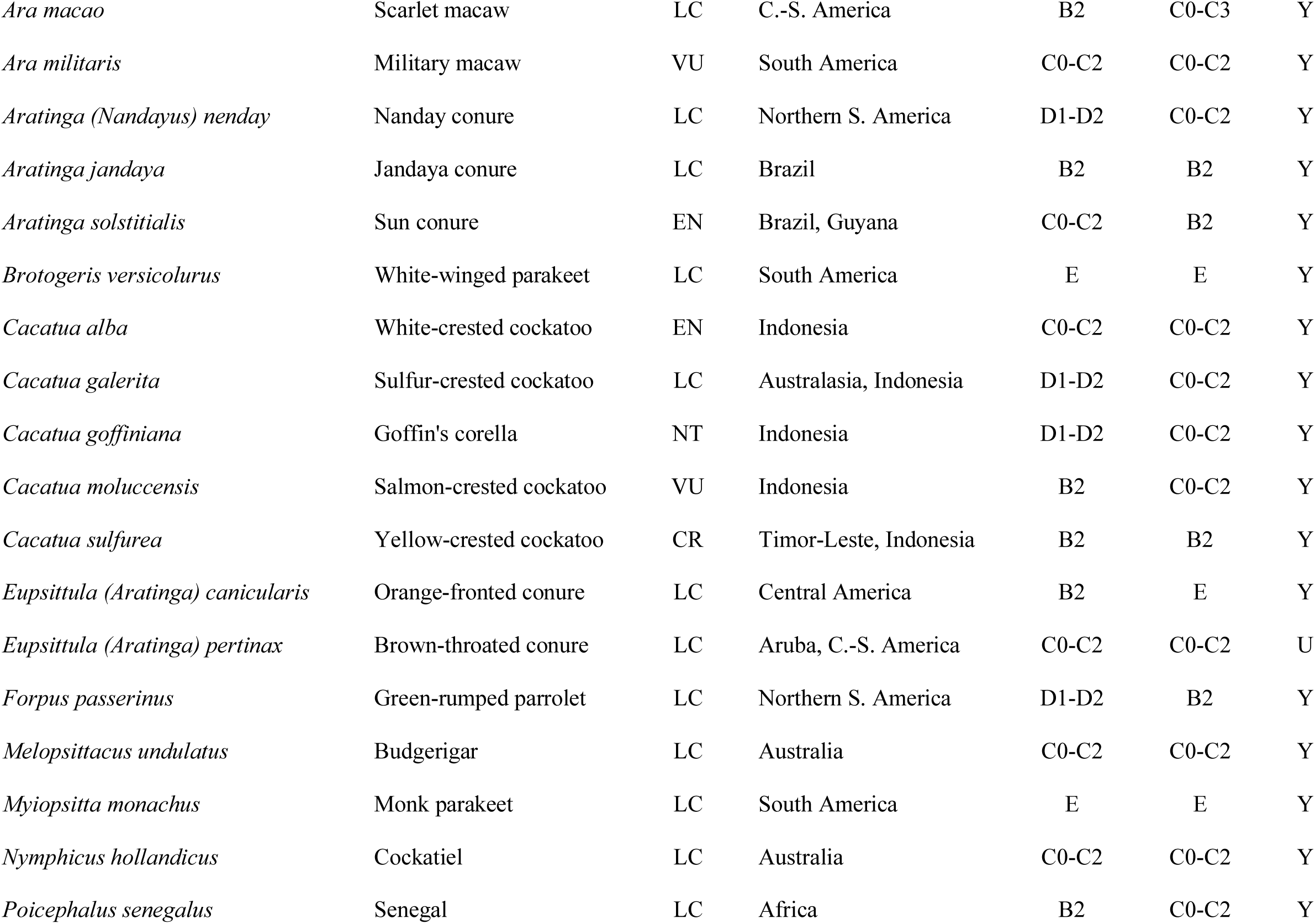
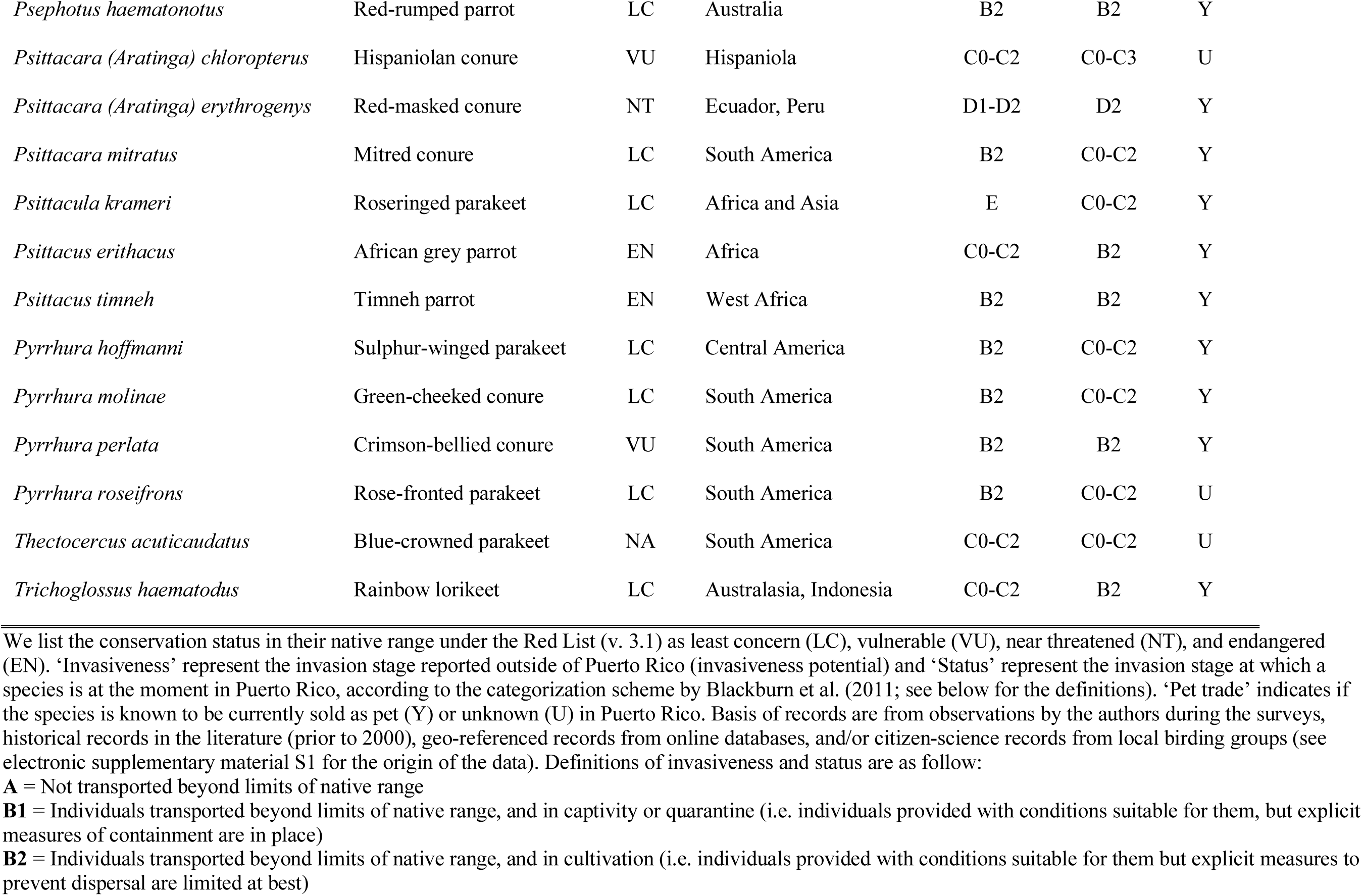
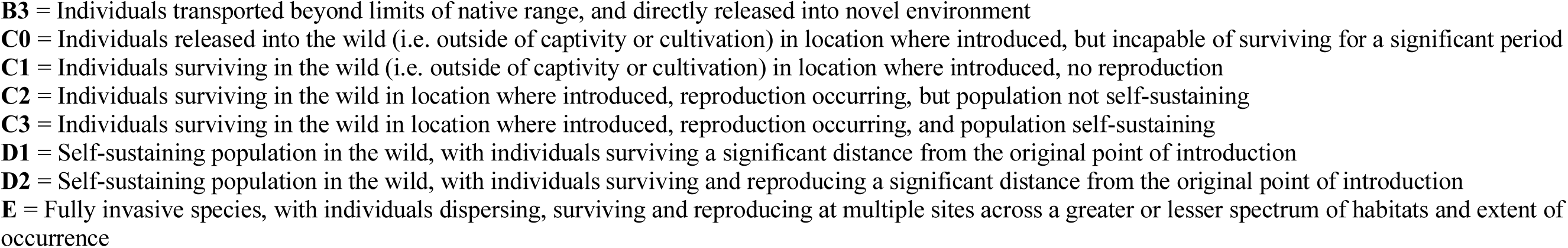
Introduced Psittaciformes reported in Puerto Rico, their native range, invasiveness and their current status on the island.

#### Sighting trends of Psittaciformes in Puerto Rico

To assess the sighting trends of Psittaciformes in Puerto Rico, we used observation count reports from *Ebird* for each species (see electronic supplementary information S2 for count data). We calculated the mean number of birds counted per municipality/year (where they have been observed), and then summed the mean number of birds per municipality to obtain the island-wide counts per year. Only species with at least 20 records were included in the subsequent analysis in this paper.

#### Distribution of Psittaciformes in Puerto Rico

We assess the current distribution of the different psittacine species found in Puerto Rico by surveying *Ebird* for geo- referenced records (see electronic supplementary information S2 for location records). After identifying psittacine species whose sighting trends showed an increase, we employed niche-modeling techniques following the methodology used by Falcón, Ackerman & Daehler (2012) & Falcón et al. (2013). To construct the distribution models, we used the Maximum Entropy Method for species distribution modelling (MaxEnt ver. 3.4.1; Phillips, Anderson & Schapire, 2006; Phillips, Dudík & Schapire, 2017). MaxEnt is a learning machine method, which uses presence only data in combination with predictive variables to model a species’ geographic distribution. Several studies have demonstrated that MaxEnt performs well at predicting species geographic distributions, and it has shown better performance and predictive ability when compared to other niche-modeling techniques (e.g., Duque-Lazo et al., 2016; Bueno et al., 2017). We used climatic layers obtained from WorldClim (ver. 2) as predictive variables in our model (http://worldclim.org/version2; Fick & Hijmans, 2017). These variables are derived from monthly temperature and rainfall values that represent annual trends, seasonality and extreme or limiting environmental factors (Hijmans et al., 2005). For our models, we selected temperature seasonality (BIO4), maximum temperature of the warmest month (BIO5), minimum temperature of the coldest month (BIO6), precipitation of the wettest month (BIO13), precipitation of the driest month (BIO14) and precipitation seasonality (BIO15) as climatic variables because they represent the extreme limiting climatic factors and their variation. The calibration area was defined as the smallest rectangle that encompassed all the location records that we used for the model plus 10 km, and restricted to the subtropics in the Americas (minimum and maximum latitude −30-30°; Corlett, 2013). For presence records, we used the geo- referenced and validated locations that we obtained from the different sources listed above, after eliminating duplicates and setting a minimum distance of 1.5 km between occurrence records (to prevent model overfitting due to similar climatic conditions of adjacent points). We randomly selected 20% of the points to test the performance of the model, and performed 10 replicates. To evaluate the performance of the model, we used the AUC statistics (area under the receiver operating characteristic curve), which provides a single measure of the model performance (Phillips, Anderson & Schapire, 2006). Models that have an excellent predictive performance have AUC values > 0.90, 0.80–0.90 are considered to have good predictive performance, while models with AUC values < 0.70 are considered poor (Swets, 1988;Manel, Williams & Ormerod, 2001; Franklin, 2010). To determine the presence-absence threshold, we used the Maximum Training Specificity plus Sensitivity threshold, which minimizes the mean of the error rate for positive and negative observations (Manel, Williams & Ormerod, 2001; Freeman & Moisen, 2008), and performs better than other thresholds in providing accurate presence predictions (Liu et al., 2005; Jiménez-Valverde & Lobo, 2007; Freeman & Moisen, 2008).

### *Ecology* of Brotogeris versicolurus *in Puerto Rico*

*Brotogeris* (Psittacidae) is a small genus (7–8 spp.) of highly abundant Neotropical parakeet species (Juniper & Parr, 1998; Forshaw, 2010), occupying varied habitats from dry savannas to tropical rain forests (del Hoyo, Elliott & Sargatal, 1997). Some of these species, if not all, have been harvested to supply the demands of the pet trade market, which is likely to have resulted in population reduction in their native habitat, while some of these species have had their populations reduced as a consequence of habitat destruction (e.g., *B. phyrrhoptera*; Birdlife International, 2016).

The white-winged parakeet is considered a widespread species in its native range of South America, occupying French Guiana, the Amazon basin from the north of Brazil to the south-east of Colombia, east of Ecuador, north of Argentina and Paraguay and the south-east of Brazil (Forshaw, 1973). Its non-native range includes Ecuador (west of Los Andes, Freile et al., 2012) California (Arrowood, 1981), and Florida (Juniper & Parr, 1998; Forshaw, 2010). Moreover, in 1969, Kepler made the first report of feral white-winged parakeets in Puerto Rico, in the area of Luquillo (Bond, 1971), and since, others have reported its presence and persistence (Pérez-Rivera et al., 1985; Raffaele, 1989; Camacho-Rodríguez, Chabert-Llompart & López-Flores, 1999). This parakeet is considered mainly a frugivorous species, although they may also use nectar and seeds as part of their diet (Janzen, 1981; Pérez-Rivera et al., 1985; Francisco, Lunardi & Galetti, 2002).

Many species of the Psittacidae family, including *B. versicolurus* roosts communally (Juniper & Parr, 1998; Brightsmith et al., 2017). The white-winged parakeet nests in arboreal termite mounds and sometimes in tree cavities (Pérez-Rivera et al., 1985). Moreover, we observed that in Puerto Rico, a proportion of the roosting population presumably break up into pairs and breed from late December to June and then eventually join the non-breeding, roosting proportion of the population. This was confirmed by Tossas, Colón & Sanders (2012) in a population in San Germán, and has also been reported in the native range (Forshaw, 1973; Juniper & Parr, 1998). In the continental U.S., fledglings have been reported to stay with the parents and return to the roosting sites after the breeding season (Schroads, 1974; Arrowood, 1981). In California, the parakeets have shown roosting site fidelity (Arrowood, 1981), and in Florida, seasonal and annual shifts in roost sites have been reported (Schroads, 1974). The birds were not marked in any of the studies, so it is impossible to determine if this involves the same individuals across time (Brightsmith et al., 2017). During the study period, we recorded observations on natural history information about the food and nest resources (plant species) used by the white-winged parakeet, and also extracted information from Pérez-Rivera et al. (1985).

To estimate the roosting population size, we used the Roost Count Method (Chapman, Chapman & Lefebvre, 1989; Gnam & Burchsted, 1991; Pithon & Dytham, 1999; Tossas, Colón & Sanders, 2012). We identified roosting sites for each population during the non-breeding season, following flocks before dusk. Then we identified areas from which the birds could be readily observed entering or leaving the roosting area. The monitoring period started from dawn in Guaynabo and before dusk in San Germán, and stopped after 20 minutes with no birds departing or entering the roost area. Since the birds gather in flocks and fly in different directions from or to the roosting area, each observer counted birds flying out of or into the roosting area in different directions to avoid double counting. If any flock returned to- (San Patricio) or left the- (San Germán) roosting area it was subtracted from the count values, as they would ultimately depart from the area again, thus avoiding double counting. We used SLR cameras with telephoto lens (70-300mm) to photograph the flocks and then count the quantity of individuals for large flocks (>16 individuals) from the photos. Days with bad weather which resulted in difficulty in counting the birds were avoided and counts were repeated if necessary. Non-breeding counts (representing the minimum population size) in Guaynabo were conducted in September of each year (2007–2009, but on 2010 it was conducted in November due to rains), and the counts in San Germán were conducted in October of each year (2007–2010). In 2010, we performed breeding counts during late March (mid-late breeding season) in both roosting populations to estimate the proportion of breeding birds.

The Roost Count Method was evaluated along with Point Transects, Line Transects and mark-recapture, using the green-rumped parrolets *(Forpus passerines)*, and the four methods produced 95% confidence intervals that overlapped and similar population estimates (Casagrande & Beissinger, 1997). To test the effectiveness of this method with the white-winged parakeet, we conducted three counts spaced a week between them from September to October 2007 in the Guaynabo population, and the population estimates exhibited low variability (range: 1,164–1,299 ±72 SD).

For the population estimates, and to calculate the population growth rates, we assumed that 1) all the parakeets roosted communally during de non-breeding season and that during the breeding season, only non-reproductive parakeets roosted communally, 2) the parakeets returned to roost in the same area after the breeding season, 3) the counts that were performed during the non-breeding season reflected the population size after mortality and emigration events shortly after the breeding season (these cannot be separated with this method), and 4) the number of parakeets counted in each roost is accurate, and 5) the estimated population size represents the minimum number of parakeets in the roost.

We calculated the population growth rates using the equation for exponential growth in populations with overlapping generations (Rockwood, 2006) re-arranged as *r* = [ln (*N_t_*/*N_0_*)]/*t* to get the *r* values, where:

*r* is the intrinsic rate of increase;

*N_t_* is the population size at time *t;*

*N_0_* is the initial population size;

*t* is the time in years

This model assumes that 1) the population has overlapping generations and that 2) the population exhibits a stable age structure. In this model, the population grows if *r* > 0, decreases if *r* < 0 and is stationary if *r* = 0. We then converted the *r* values to the finite rate of increase per yr, λ, with the equation λ, = *e^r^*. When λ < 1 the population decreases, the population is stable if λ = 1 and the population increases if λ > 1.

To determine the proportion of the breeding population, we subtracted the number of parakeets counted during the breeding censuses of 2010 from the population estimates of the 2009 non-breeding censuses. Since both adults have been reported to roost in the nest until all the chicks are fledged (Schroads, 1974), we divided the total number of breeding individuals by two to get the approximate number of breeding pairs.

### Statistical analyses and data visualization

We performed all data pre-processing, obtained summary statistics, and visualizations using R ver. 3.3.3 (R Core Team, 2017), and packages ‘ggplot2’ (Wickham, 2016), ‘raster’(Hijmans, 2017), ‘zoo’ (Zeileis & Grothendieck, 2005), ‘plyr’ (Wickham, 2011), and the output from MaxEnt. We mapped occurrence records and visualized distribution results using QGIS ver. 2.18.14-Las Palmas de G. C. (QGIS Development Team, 2017).

## Results

### Introductions of Psittaciformes to Puerto Rico

#### Historical and current status of Psittaciformes

We found historical records for 18 species of Psittaciformes reported by the year 2000, with eight of those breeding. On the other hand, we found that at least 46 species of Psittaciformes are currently present on the island (Fig. 1; Table 1), of which 24% are only found in the pet trade, 48% have been observed in the wild (present), but are probably not established, and 28% are established and known to have been or be currently breeding (Table 1). At least 85% of the species are currently available for sale in the pet trade. Of the 46 species of Psittaciformes found in Puerto Rico, at least 63% have been reported in the wild elsewhere (but it is unknown whether they are breeding), and 26% are considered as established or invasive (i.e., breeding outside their native range and expanding their territory, but not necessarily causing negative impacts).

**Figure 1:**
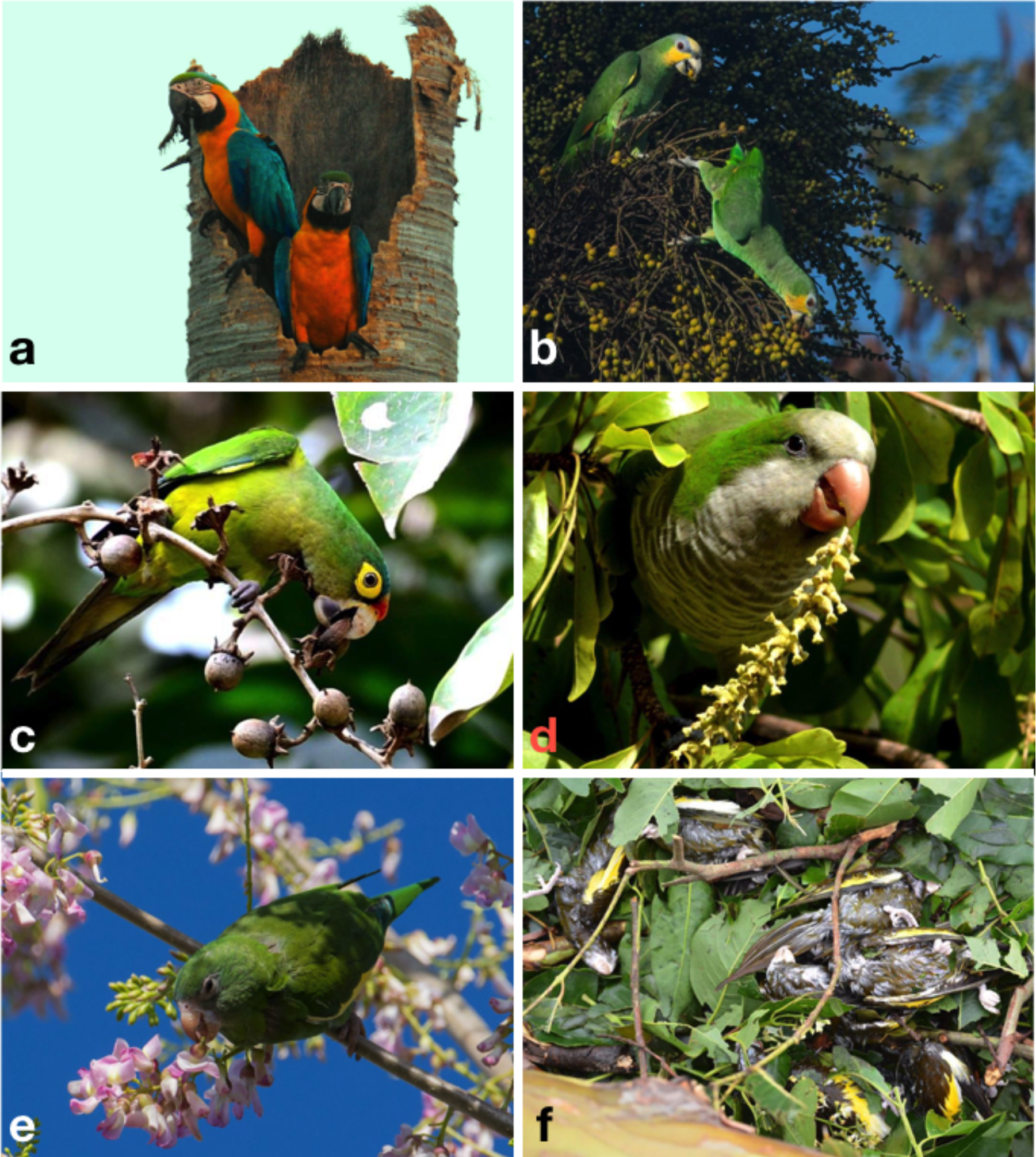
Some of the species of Psittaciformes that occur in the wild in Puerto Rico, and Hurricane Maria-related mortality. A pair of blue-and-yellow macaws (*Ara ararauna*) in their nest on *Roystonea borinquena* (Aracaceae; a), an orange-winged amazon (*Amazona amazonica*) eating palm fruits (Aracaceae; b), an orange-fronted parakeet (*Eupsittula canicularis*) foraging on flowers (Fabaceae; c), a monk parakeet (*Myiopsitta monachus*) eating the flower buds of *Bucida buceras* (Combretaceae; d), a white-winged parakeet (*Brotogeris versicolurus*) eating flower buds (Fabaceae; e), and six out of dozens of white-winged parakeets that died during Hurricane María in 2017 (f). Photo credits: Yoly Pereira (a), Julio Salgado (b, e), Kurt Miller (c), Sonia Longoria (d), Dinath Fourquet (f).

#### Sighting trends of Psittaciformes in Puerto Rico

We found sufficient reports to calculate the island-wide sighting trends for 10 species of Psittaciformes in Puerto Rico (Fig. 2–3). Four species exhibited population increase, three species showed stable populations, and three species exhibited population decrease. For the rest of the species, the count numbers were too low and/or temporal resolution was too short to calculate trends. Of the species with population increase, the white-winged parakeet showed the largest population increase (Fig. 3), followed by the monk parakeet (*Myiopsitta monachus*), the red-masked conure (*Psittacara. (Aratinga) erythrogenys*) and the orange-fronted parakeet (*Eupsittula canicularis*; Fig. 2). The orange winged amazon (*Amazona amazonica*) showed a population increase and later stabilized, while the white-crested cockatoo (*Cacatua alba*) showed a stable population trend, albeit with low numbers. The blue-and-yellow macaw (*Ara aranaura*) exhibited a decrease from 1996 to 2002 (with up to 22 individuals reported) in the Metropolitan Area of San Juan, and later recovered and stabilized with about 14 individuals. The green-checked amazon (*Amazona viridigenalis*) showed a steep decrease after reaching a total of 53 individuals per mean count/municipality/year, with low counts after 2006. Similar trends were observed in the white-fronted amazon (*A. albifrons*), but in much lower numbers. Finally, the rose-ringed parakeet (*Psittacula krameri*) showed a sustained decrease after a maximum of 11 individuals were reported in 2011.

**Figure 2:**
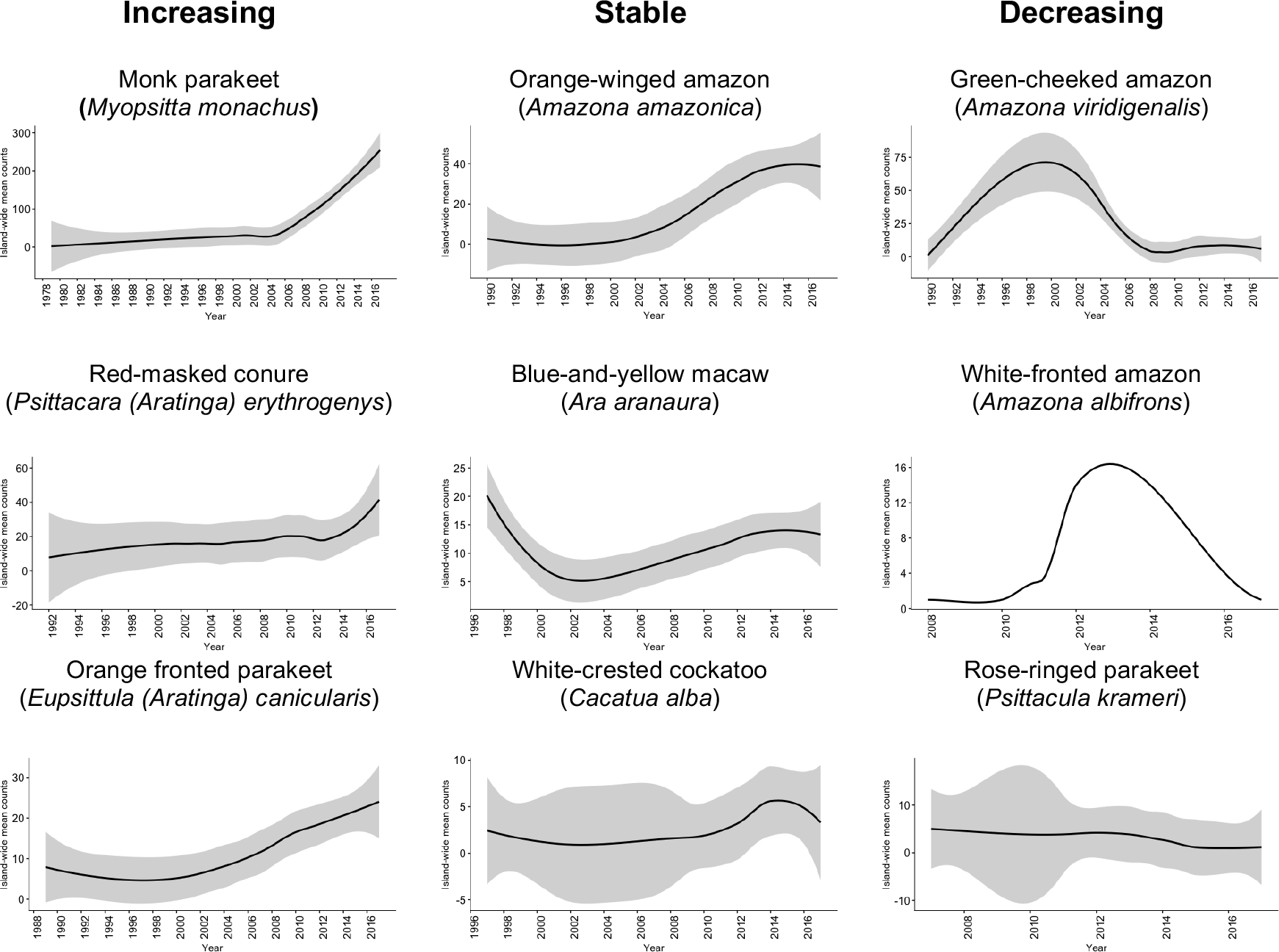
Sighting trends of different species of Psittaciformes in Puerto Rico showing population increase, stable populations and population decrease. Island-wide sighting trends were calculated as the sum of the mean number of birds counted per year/municipality (data from Ebird). Grey shading indicates the 95% CI based on the local weighted scatterplot smoothing (loess).

**Figure 3:**
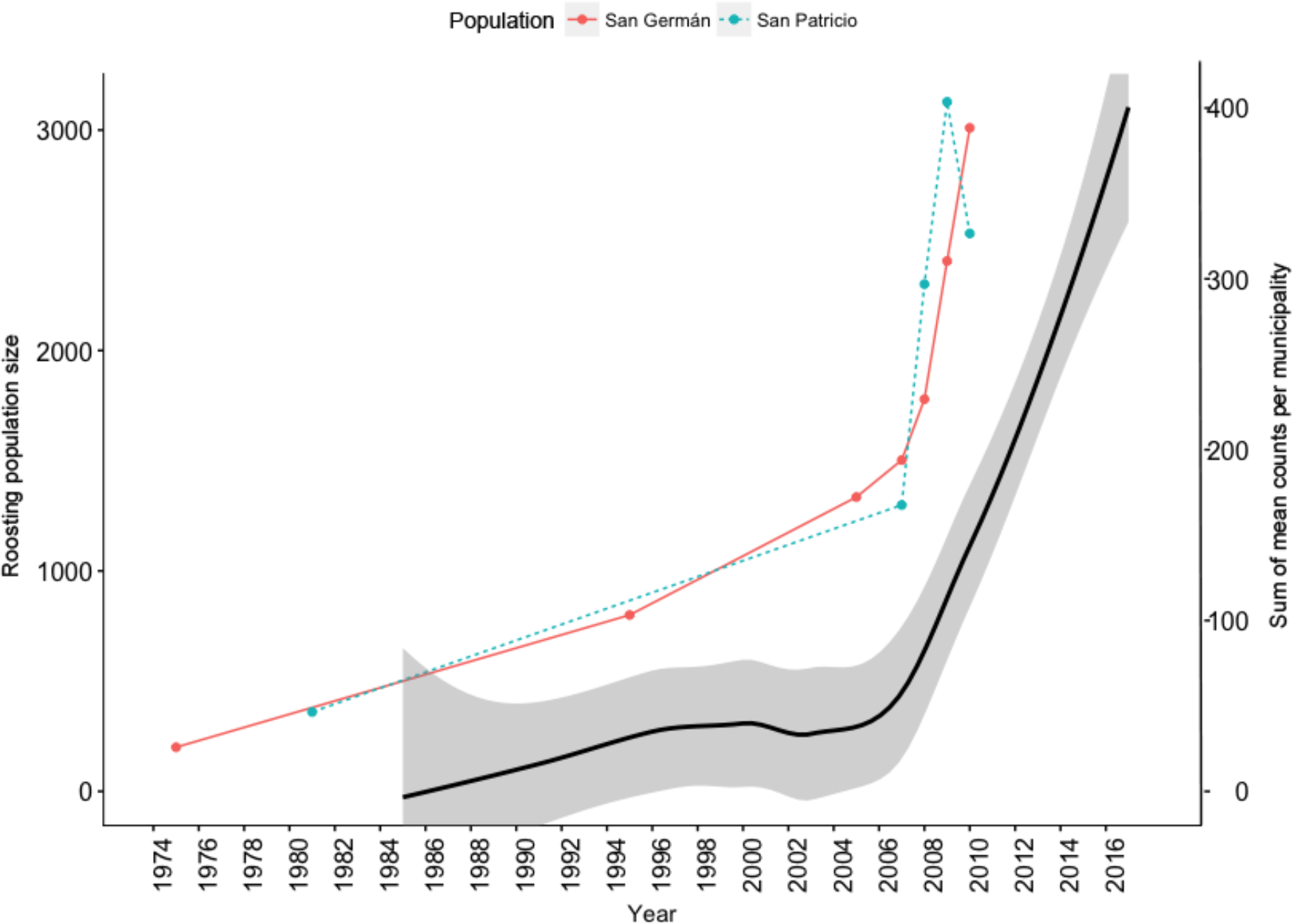
Island-wide population increase of the white-winged parakeet (*Brotogeris versicolurus*) in Puerto Rico (black line) and population increase in two populations (roosting sites) over the years. Island-wide population increase was calculated as the sum of the mean number of birds counted per year/municipality (data from Ebird). Grey shading indicates the 95% CI based on the local weighted scatterplot smoothing (loess).

#### Distribution of Psittaciformes in Puerto Rico

Overall, we obtained 6,905 locality records for 26 species of Psittaciformes in the wild that spanned from 1960 to 2017 (in *Ebird*; see electronic supplementary information S1 for detailed species-specific results and S2 for location records). Moreover, we obtained 279 sighting records for 10 species from local groups (see electronic supplementary information S3 for links to species-specific reports). Sightings and reports of Psittaciformes in Puerto Rico practically cover the whole island, and are especially dense in coastal and highly (human) populated areas (Fig. 4), but the geographic extent of the distribution varies by species (see electronic supplementary information S4 for the distribution of locations for each species reported in the island).

**Figure 4:**
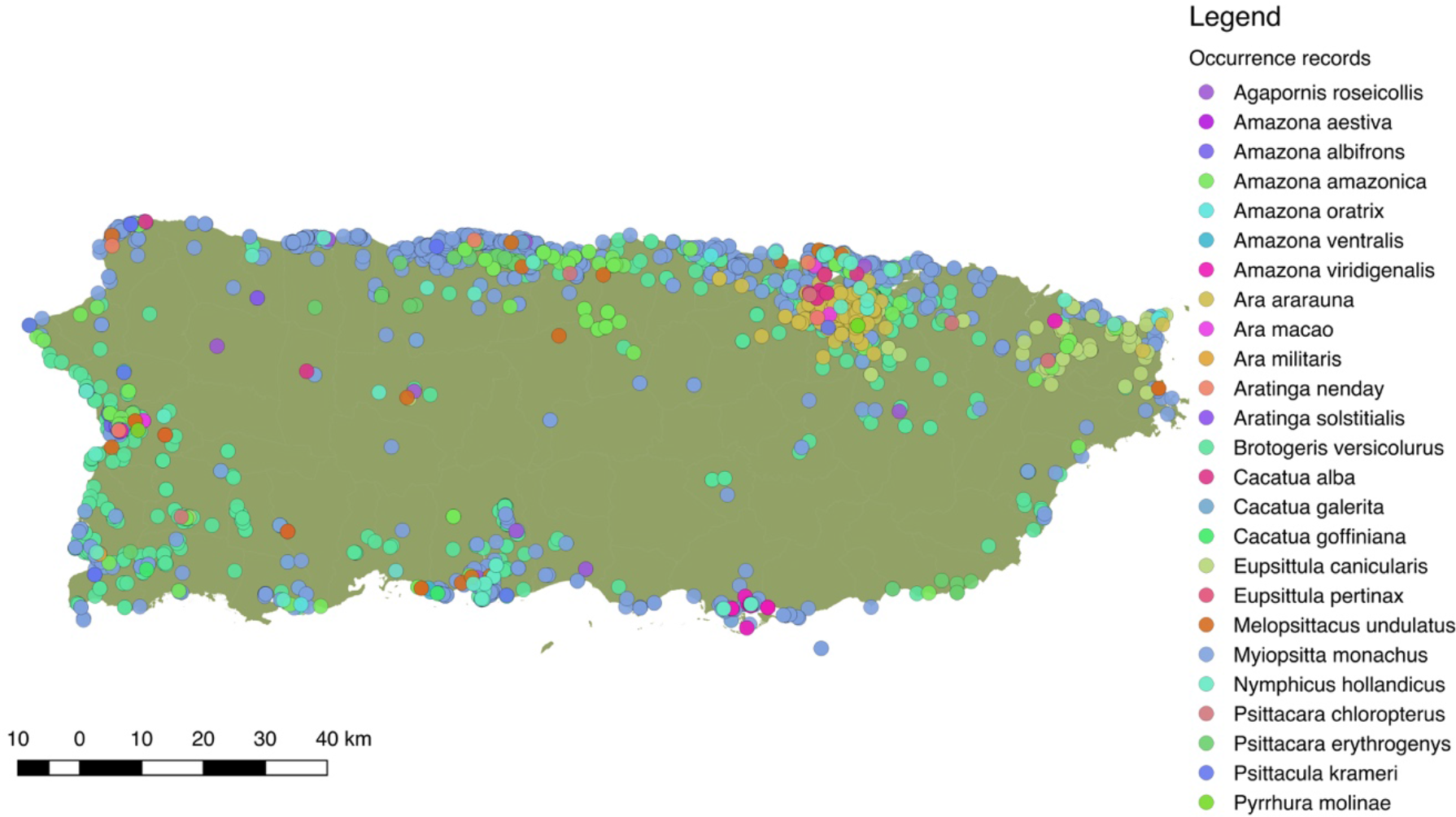
Distribution of 25 species of Psittaciformes in Puerto Rico, depicted by the different colors. Records originated from observations made by the authors, online databases, and reports from local birding groups (see methods).

More specifically, for the white-winged parakeet, the different locality record sources yielded a total of 2,520 occurrence records in Puerto Rico. Historic reports on the distribution of *B. versicolurus* show that parakeets were present in Luquillo (east) in small numbers by the 1960’s (Kepler *in* Bond 1971). Moreover, a small population was found breeding in the municipality of Naguabo (east) and about 360 individuals were reported in Guaynabo by 1985, were the San Patricio population is presently located (Fig. 5; Pérez-Rivera et al., 1985). Furthermore, the only other historic record are from the population in San Germán, which was estimated at 800 individuals by 1995 (Camacho-Rodríguez, Chabert-Llompart & López-Flores, 1999). Since then, the parakeet populations have expanded significantly throughout the island, especially to coastal and heavily (human) populated zones, but they have also been occasionally observed in the central mountainous regions (Fig. 5).

**Figure 5:**
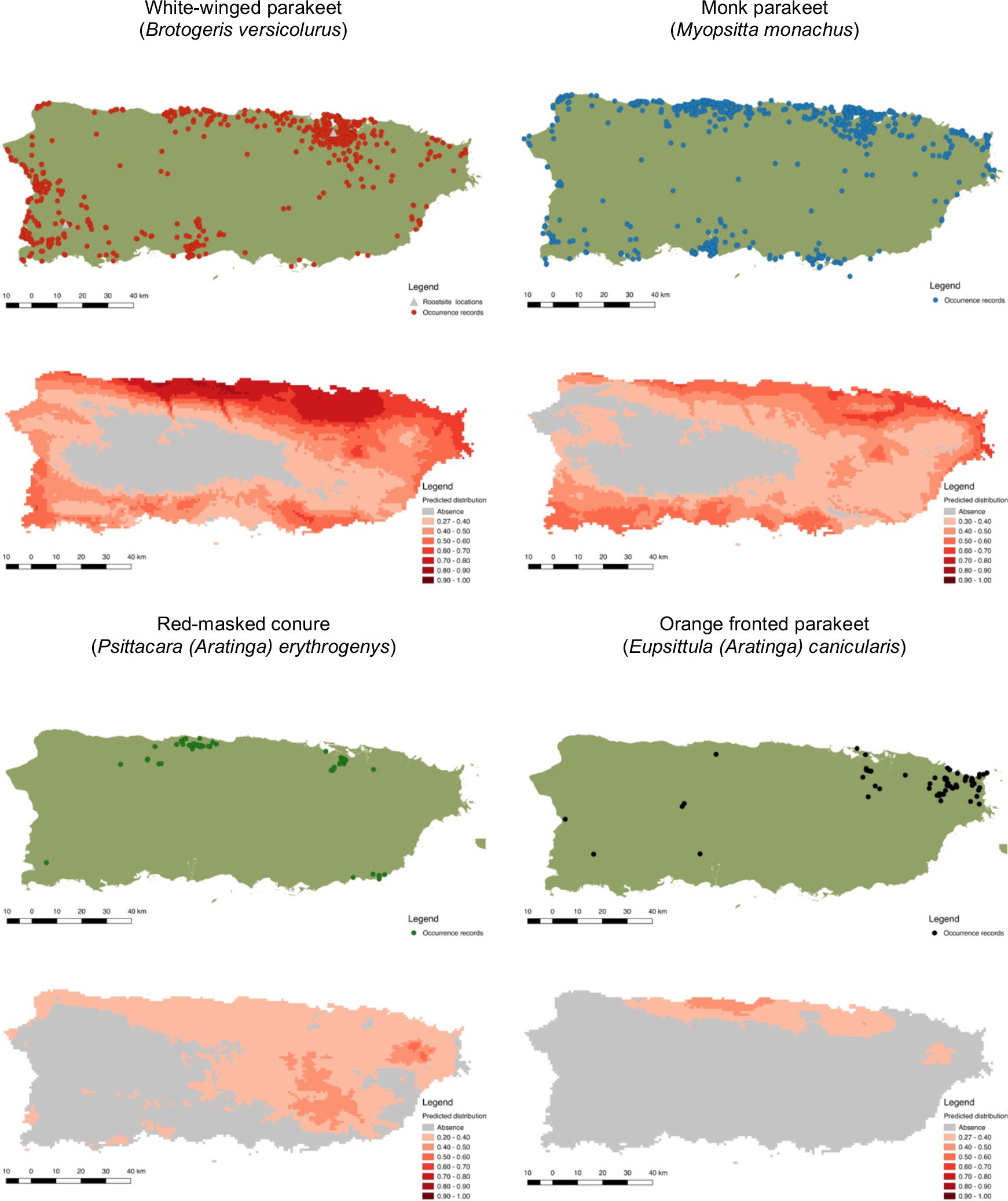
Current distribution of Psittaciformes in Puerto Rico whose populations are increasing (based on sighting trends, top figures with points) and the predicted distribution of the species based on the maximum entropy model (MaxEnt, bottom figures with continuous color). Warmer colors depict higher suitability.

For predicting the potential distribution of the psittacine species whose populations are increasing in Puerto Rico, we obtained a total of occurrence 106,493 records from their native range and from their invasive range (including Puerto Rico; see electronic supplementary information S2). Our models performed good to excellent, with test AUC values ranging from 0.82–0.94 (Table 2), and the presence records for the four species are within the predicted distribution in its native range, indicating a good model fit. Furthermore, the models predicted suitable areas for all four species in Puerto Rico (Fig.5).

**Table 2:** Occurrence locations and MaxEnt summary statistics for predicting the distribution of four species of Psittaciformes in Puerto Rico.

| Species | Occ. loc. | Training AUC | Test AUC | MS+ST |
| --- | --- | --- | --- | --- |
| *Brotogeris versicolurus* | 1,119 (557) | 0.91 ($\pm 0.001$) | 0.92 ($\pm 0.006$) | 0.27 ($\pm 0.024$) |
| *Myopsitta monachus* | 15,985 (707) | 0.82 ($\pm 0.001$) | 0.82 ($\pm 0.004$) | 0.30 ($\pm 0.031$) |
| *Psittacara erythrogenys* | 1,392 (42) | 0.94 ($\pm 0.002$) | 0.94 ($\pm 0.003$) | 0.20 ($\pm 0.036$) |
| *Eupsittula canicularis* | 3,707 (55) | 0.84 ($\pm 0.001$) | 0.84 ($\pm 0.005$) | 0.27 ($\pm 0.014$) |

Total occurrence locations used for the model (Occ. loc.), and the subset of occurrence locations for Puerto Rico given in parentheses. Model performance for predicting the distribution of four species of Psittaciformes in Puerto Rico given by the training (80% occurrence points) and test (20% occurrence points) Area Under the ROC Curve (AUC), and the Maximum training Sensitivity plus Specificity presence-absence threshold (MS+ST). Values given are the mean ( ± SD) based on ten replicates.

The predicted distribution for the white-winged parakeet in Puerto Rico showed the highest suitable areas in the north-central part of the island, but also included the other coastal areas. Only in the central and central-west part of the island, where the Central Cordillera occurs, did MaxEnt predicted unsuitable areas for the species. In general, *B. versicolurus* occupies virtually all the areas with the highest suitability. Similar areas were predicted with suitable climatic conditions for the monk parakeet, although there are areas in the west of the island predicted as unsuitable where the monk parakeet has been reported. The models also predicted suitable areas for the red-masked conure and the orange-fronted parakeet, some of which they occupy. Both species have lower predicted climatic suitability than white-winged and monk parakeets, and the orange-fronted parakeet has a smaller predicted suitable area than the other species. In general, our models predicted suitable areas outside the current range of all four species, indicating the possibility for further range expansion.

### *Ecology* of Brotogeris versicolurus *in Puerto Rico*

During the study period, we made observations on the diet of the white-winged parakeet in the Metropolitan Area of San Juan. They use mostly exotic species of plants as food resources, and as in their native range, they are primarily frugivorous, but also consume flowers, seeds and sap (Table 3; complimented with data from Pérez-Rivera et al., 1985). The annual breeding cycle of the white-winged parakeet in the Metropolitan Area of San Juan starts in January, at the end of the dry season, with reproductive pairs starting to inspect possible nests, and continues through February with pairs constructing their nests. They may use new termite mounds, or re-use termite mounds used in previous years for nesting. Between February and April, reproductive pairs nest and raise their chicks while non-reproductive parakeets presumably stay at the roosting sites. Most chicks are fledged by June, and presumably join the roosting parakeets along with their parents. In terms of nesting, although there were tree cavities likely suitable for the white-winged parakeets in some places where nesting occurred, all nest we found were constructed in termite mounds of *Nasutitermes sp.* in native and exotic trees (Table 3). Two nests inspected with a pipe inspection camera revealed one clutch of three eggs and one clutch of four eggs. Moreover, five nests inspected with the same method revealed a mean of 2.14 (±0.69) chicks per nest. Furthermore, upon inspection of a chick that fell from another nest, we found feather mites (Acarina), likely of the genus *Pararalichus.*

**Table 3:**
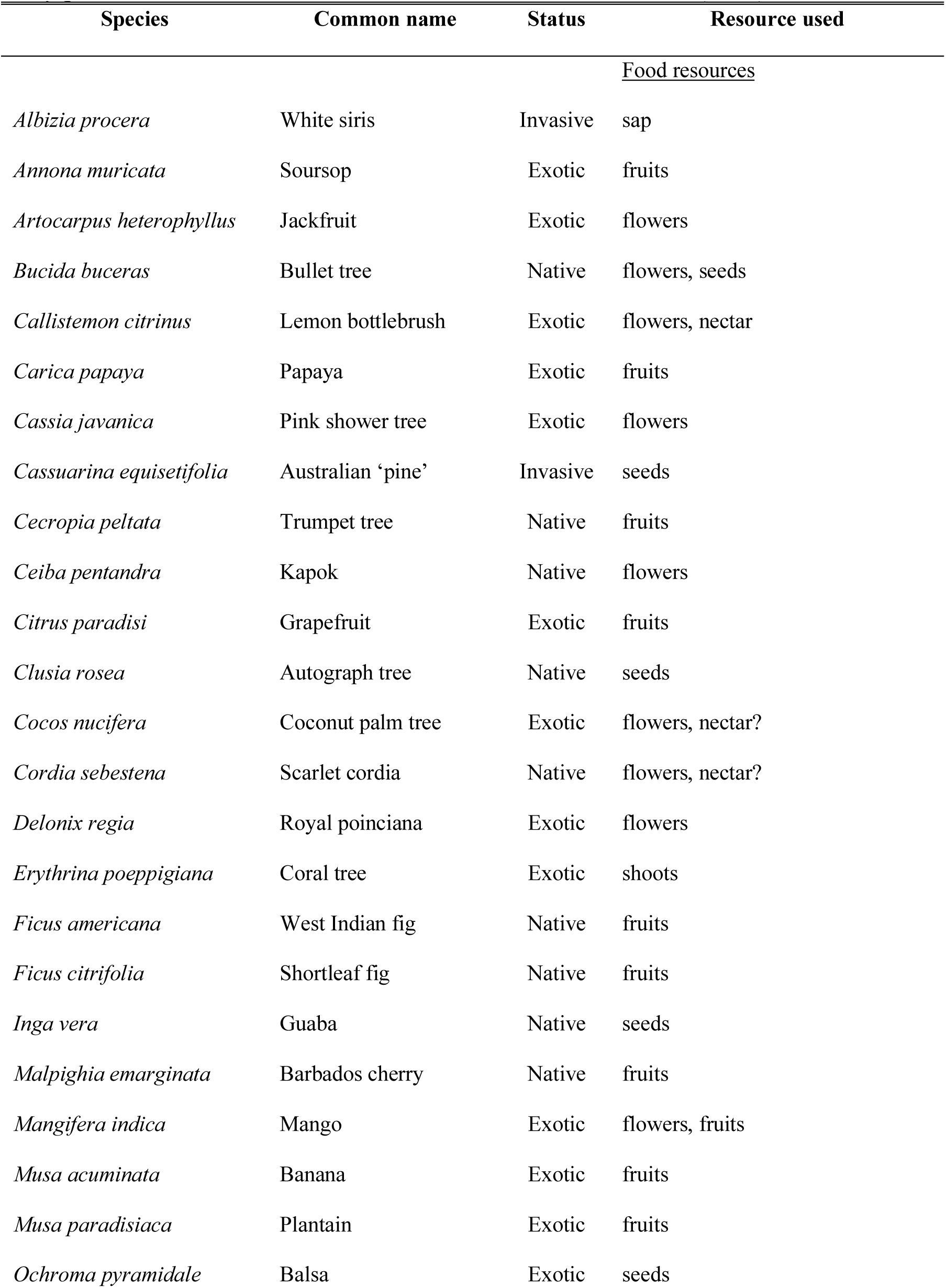
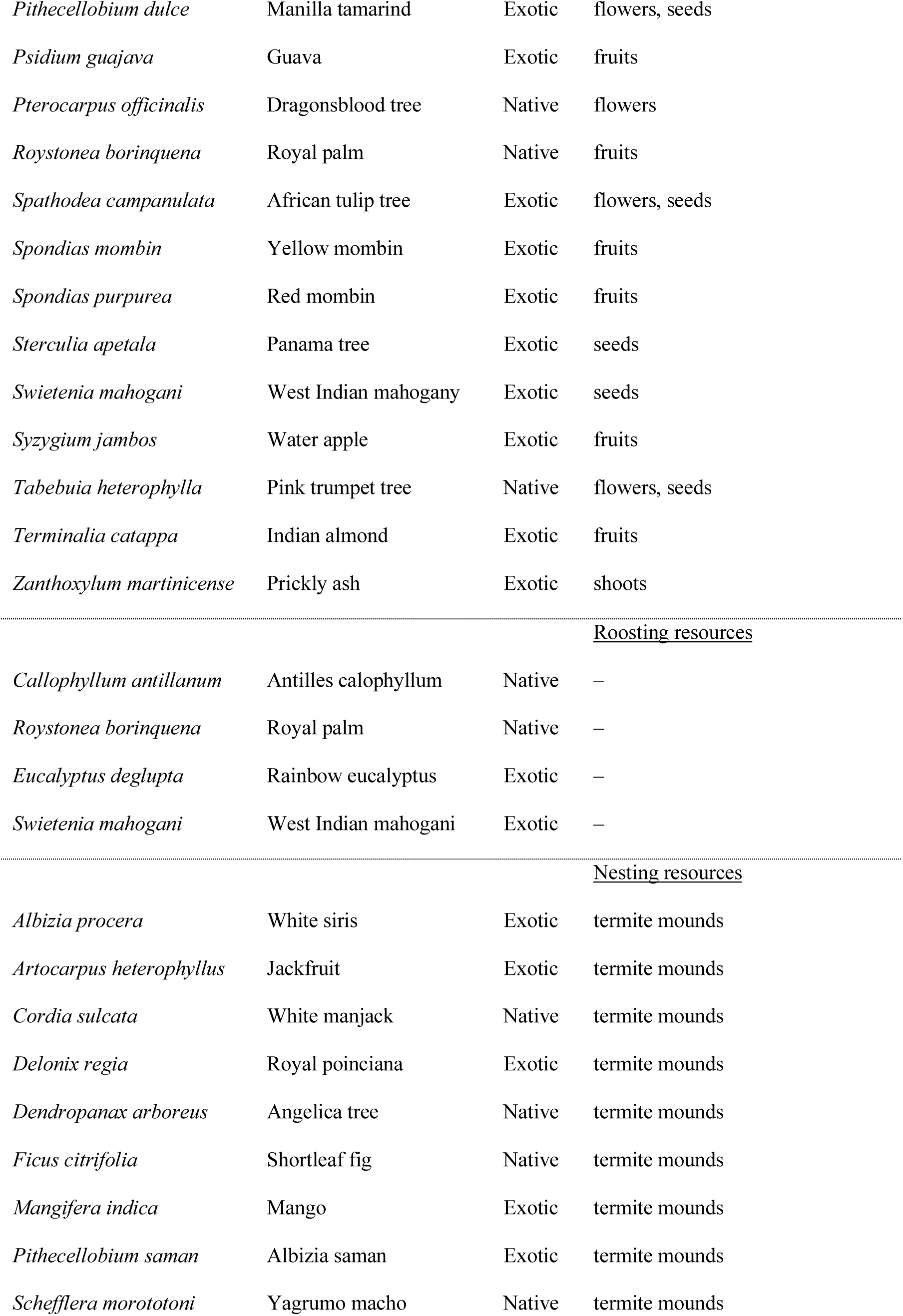
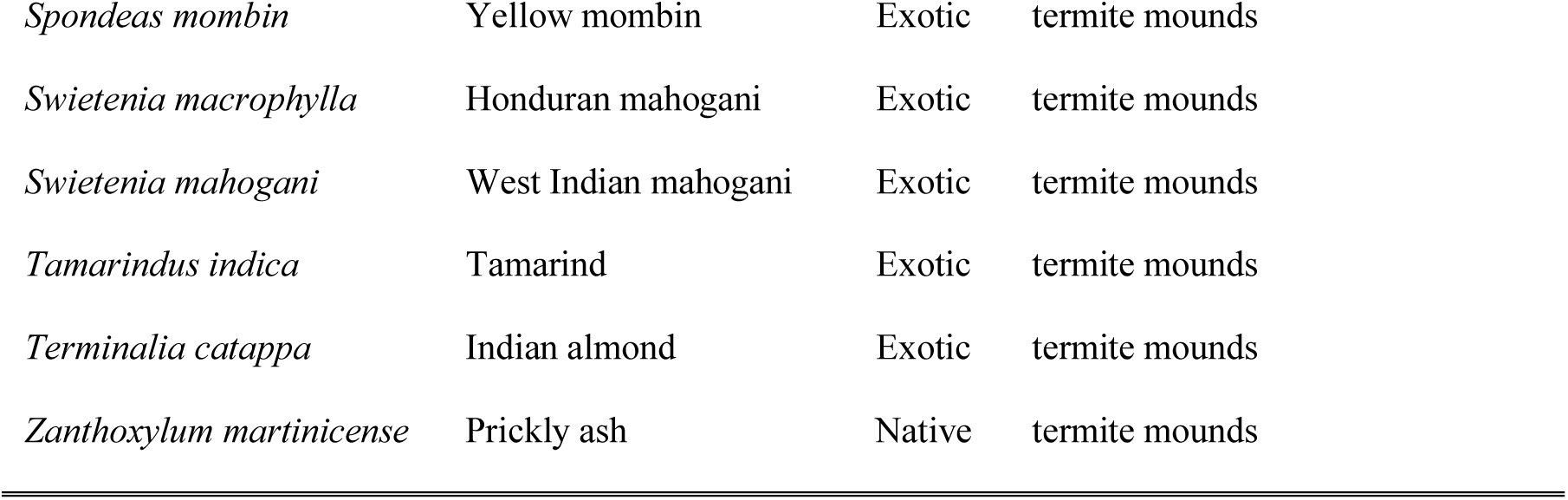
Plant resources used by the white-winged parakeet, *Brotogeris versicolurus* for food and nesting in Puerto Rico. Information based on observations made during the study period, with additional information from Pérez-Rivera et al. (1985).

| Species | Common name | Status | Resource used |
| --- | --- | --- | --- |
|  |  |  | <u>Food resources</u> |
| *Albizia procera* | White siris | Invasive | sap |
| *Annona muricata* | Soursop | Exotic | fruits |
| *Artocarpus heterophyllus* | Jackfruit | Exotic | flowers |
| *Bucida buceras* | Bullet tree | Native | flowers, seeds |
| *Callistemon citrinus* | Lemon bottlebrush | Exotic | flowers, nectar |
| *Carica papaya* | Papaya | Exotic | fruits |
| *Cassia javanica* | Pink shower tree | Exotic | flowers |
| *Cassuarina equisetifolia* | Australian ‘pine’ | Invasive | seeds |
| *Cecropia peltata* | Trumpet tree | Native | fruits |
| *Ceiba pentandra* | Kapok | Native | flowers |
| *Citrus paradisi* | Grapefruit | Exotic | fruits |
| *Clusia rosea* | Autograph tree | Native | seeds |
| *Cocos nucifera* | Coconut palm tree | Exotic | flowers, nectar? |
| *Cordia sebestena* | Scarlet cordia | Native | flowers, nectar? |
| *Delonix regia* | Royal poinciana | Exotic | flowers |
| *Erythrina poeppigiana* | Coral tree | Exotic | shoots |
| *Ficus americana* | West Indian fig | Native | fruits |
| *Ficus citrifolia* | Shortleaf fig | Native | fruits |
| *Inga vera* | Guaba | Native | seeds |
| *Malpighia emarginata* | Barbados cherry | Native | fruits |
| *Mangifera indica* | Mango | Exotic | flowers, fruits |
| *Musa acuminata* | Banana | Exotic | fruits |
| *Musa paradisiaca* | Plantain | Exotic | fruits |
| *Ochroma pyramidale* | Balsa | Exotic | seeds |
| *Pithecellobium dulce* | Manilla tamarind | Exotic | flowers, seeds |
| *Psidium guajava* | Guava | Exotic | fruits |
| *Pterocarpus officinalis* | Dragonsblood tree | Native | flowers |
| *Roystonea borinquena* | Royal palm | Native | fruits |
| *Spathodea campanulata* | African tulip tree | Exotic | flowers, seeds |
| *Spondias mombin* | Yellow mombin | Exotic | fruits |
| *Spondias purpurea* | Red mombin | Exotic | fruits |
| *Sterculia apetala* | Panama tree | Exotic | seeds |
| *Swietenia mahogani* | West Indian mahogany | Exotic | seeds |
| *Syzygium jambos* | Water apple | Exotic | fruits |
| *Tabebuia heterophylla* | Pink trumpet tree | Native | flowers, seeds |
| *Terminalia catappa* | Indian almond | Exotic | fruits |
| *Zanthoxylum martinicense* | Prickly ash | Exotic | shoots |
| Roosting resources |  |  |  |
| *Callophyllum antillanum* | Antilles callophyllum | Native | — |
| *Roystonea borinquena* | Royal palm | Native | — |
| *Eucalyptus deglupta* | Rainbow eucalyptus | Exotic | — |
| *Swietenia mahogani* | West Indian mahogani | Exotic | — |
| Nesting resources |  |  |  |
| *Albizia procera* | White siris | Exotic | termite mounds |
| *Artocarpus heterophyllus* | Jackfruit | Exotic | termite mounds |
| *Cordia sulcata* | White manjack | Native | termite mounds |
| *Delonix regia* | Royal poinciana | Exotic | termite mounds |
| *Dendropanax arboreus* | Angelica tree | Native | termite mounds |
| *Ficus citrifolia* | Shortleaf fig | Native | termite mounds |
| *Mangifera indica* | Mango | Exotic | termite mounds |
| *Pithecellobium saman* | Albizia saman | Exotic | termite mounds |
| *Schefflera morototoni* | Yagrumo macho | Native | termite mounds |
| *Spondeas mombin* | Yellow mombin | Exotic | termite mounds |
| *Swietenia macrophylla* | Honduran mahogani | Exotic | termite mounds |
| *Swietenia mahogani* | West Indian mahogani | Exotic | termite mounds |
| *Tamarindus indica* | Tamarind | Exotic | termite mounds |
| *Terminalia catappa* | Indian almond | Exotic | termite mounds |
| *Zanthoxylum martinicense* | Prickly ash | Native | termite mounds |

The San Germán population of white-winged parakeets increased from 1,503 birds in 2007 to 3,010 birds in 2010. However, although the San Patricio population was steadily increasing from 2007–2009, from 1,299 to 3,128 individuals, a population decline occurred in 2010, and the population was reduced to 2,531 individuals (Fig. 3). Both populations exhibited overall positive population growth rates, with a geometric mean λ = 1.24 (±0.48) in San Patricio, and λ, = 1.26 (±0.08) in San Germán during the study period (Table 4). If we consider the population estimates prior and after to our population estimates (a period of 29 yr in San Patricio, and 40 yr in San Germán), both populations exhibited *λs* of 1.20 and 1.17, respectively. Moreover, the island-wide sum of the mean counts per municipality shows an increase of birds from 9 parakeets in 1985 to 375 parakeets in 2017 among 55 of the 78 municipalities in Puerto Rico (Fig. 3). Based on the breeding census, we estimated that 40% of the population at San Patricio was breeding (625 pairs), and 42% of the population at San Germán (512 pairs). Based on the number of birds retuning to the roost after the breeding season, the number of fledglings (recruitment) was estimated in 604 for San Germán (assuming 2 chicks per pair), but because of the population decline, it was not possible to estimate this for the San Patricio population.

**Table 4:**
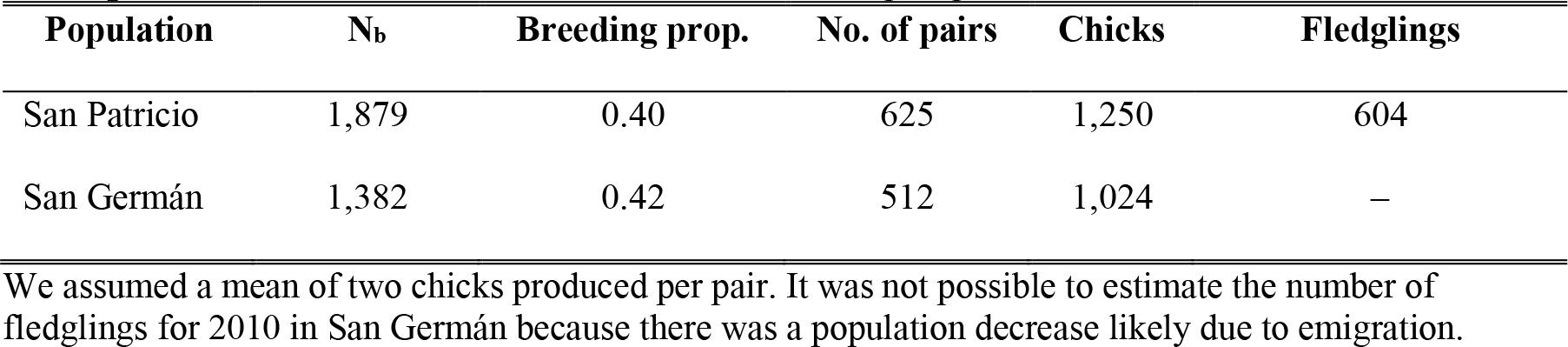
Population estimates and growth rates of the white-winged parakeet (*Brotogeris versicolurus*) at two sites in Puerto Rico. Counts were performed during the non-breeding season (N_nb_)

**Table 4:**
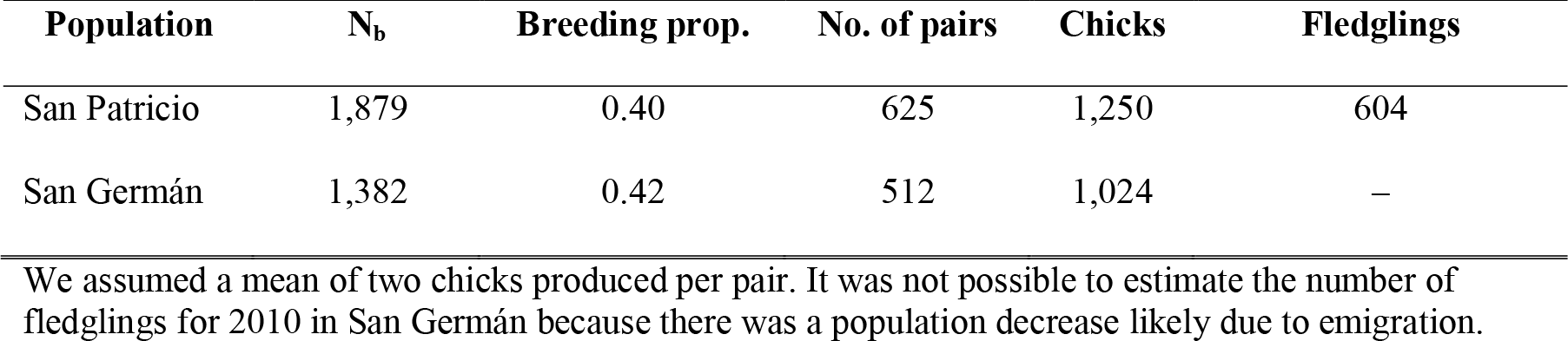
Breeding proportion of the population based on the 2010 breeding (N_b_) and non-breeding season counts at the roosting site, the estimated number of pairs, expected chick production and the estimated number of fledglings.

## Discussion

### Introductions of psittaciformes to Puerto Rico

In this study, we sought to assess the status of non-native Psittaciformes in Puerto Rico, and the invasive ecology of the white-winged parakeet. We found a total of 46 psittacine species present on the island. Of the 8 species historically reported as established (breeding) in the wild, we confirmed that six still persist, and at least nine species are currently breeding (11 if we consider all species breeding at any given time throughout the historical survey). Moreover, we found clear evidence of invasiveness for the white-winged parakeets, and sighting trends and observation suggest the same for monk parakeets. To our knowledge, Puerto Rico is the place with the highest number of wild observed and established exotic parrot species in the world based on surveys in other places where parrot populations occur (e.g., (Runde, Pitt & Foster, 2007; Mori et al., 2013; Symes, 2014).

Introduction of Psittasiformes to Puerto Rico have been solely via the pet trade. According to Forshaw (1973), the Hispaniola amazon apparently was introduced to Puerto Rico because managing agencies refused to give the importation permits for a shipment brought from the Dominican Republic by boat, which resulted in the deliberate release of ‘hundreds’ of parrots. The Hispaniola amazon is currently established and breeding in Mayagüez, but their numbers appear to be low (∼10 individuals; WF, pers. obs.). It is unknown whether the present population is the result of the release reported by Forshaw (1973) and/or the subsequent release of pet parrots into the wild. However, most Psittaciformes observed and/or established in the wild in Puerto Rico are believed to be the result of intentional or accidental releases by owners who bought them through the pet trade. For example, Pérez-Rivera (1985) made the first report of 10 individuals of the red-masked conure in Cataño. We are unaware of any existing populations of this conure in Cataño, but close-by in Río Piedras (San Juan), a population of about 20 individuals have been known since 2010 and are breeding, using cavities in buildings of the University of Puerto Rico as nests (WF pers. obs.), and there are reports of 30 individuals in the nearby Guaynabo municipality (adjacent to San Juan and Cataño municipalities). Moreover, there is a population established in the municipality of Arecibo (northcentral), with over 50 individuals reported. However, not all introduced species succeeded, for example, a ‘small group’ of masked lovebirds (*Agapornis personata*) was reported in Caguas between 1982–1984 (Pérez-Rivera 1985). No details were provided on whether the population was breeding or not. Based on the occurrence records obtained, and our observations, this species is currently absent in the wild in Puerto Rico, thus it is highly probable that this flock went extinct.

Observations on flock size and sighting trends of Psittaciformes in Puerto Rico indicate that the different species are experiencing different dynamics. For example, white-winged (discussed below in more detail) and monk parakeets are the only two species that appear to be growing exponentially. Records for the monk parakeets started in 1979 throughout coastal and densely populated (humans) areas (mainly San Juan and Ponce), and by 1987, flocks of up to 30 individuals were observed. By the late 1990’s sightings outside these two municipalities started, and became more common throughout the island as time passed probably as a result of population growth and range expansion. The orange-winged amazons exhibited an increase in sightings since the 1990’s, and by 2014, the population seemed to be stabilized. The maximum number of individuals reported is 39, but there is a population roosting in the municipality of Morovis with over 100 individuals (not included in the count data; J. Salgado Vélez, pers. comm.).

Moreover, the orange-winged amazons are the most widespread of the amazon parrots in Puerto Rico. The blue-and-yellow macaws apparently experienced a population decline in the Metropolitan Area of San Juan, and later increased and stabilized. Furthermore, a population of at least 15 individuals is breeding in the municipality of Orocovis, ∼29 km away from the other population (not included in count data), and some have been sighted in Cabo Rojo (west of the island). The sighting trends of the white-crested cockatoo seems to indicate the population is stable with low numbers (2–15 individual), and localized in the adjacent municipalities of Bayamón and Guaynabo in the Metropolitan Area. Interestingly, the green- and white-fronted amazons exhibited a sighting decline after initially increasing. The green-fronted amazon has two reported populations; one in the municipality of Mayagüez (west) with up to 10 individuals recorded, and one in the municipality of Salinas (southeast) with up to 30 individuals recorded. Similarly, the white-fronted amazon, which is restricted to Mayagüez, exhibited a reduction in sightings, from up to 11 individuals in 2011, to 1–2 individuals in recent years.

Moreover, although uncommon, there have been reports of rose-ringed parakeet in Puerto Rico, and at least 12 individuals recorded in Aguadilla in 2012. However, their sighting trends indicate a decrease in numbers, so it’s possible that the group failed to establish, and the recent reports are of escaped pets. However, rose-ringed parakeets can easily be obtained through the online pet market, and has become more popular as a pet in recent years. Therefore, more releases are possible, and the possibility of rose-ringed parakeets establishing in Puerto Rico is latent.

Psittaciformes in Puerto Rico are more commonly distributed along the coastal region, and areas with the highest occurrence records coincide with densely populated areas by humans, especially the Metropolitan Area of San Juan. However, the location and the extent of the geographic distribution of Psittaciformes varies between species. The most recorded and widespread species are those whose population trends (based on sightings) seem to be increasing, and their current distribution coincide with areas predicted to be climatically suitable. Moreover, suitable areas are present outside the areas where these different species occur, indicating the potential for continued range expansion. Of these, the white-winged and the monk parakeets are undoubtedly the most sighted and widespread species of Psittaciformes in Puerto Rico.

Perhaps the most important factor promoting the success of Psittaciformes outside their native range is the sheer number of individuals that were and are available through the pet trade. Invasion success for exotic bird species is positively influenced by the number of individuals available on the market (Carrete & Tella, 2008), and species more commonly found in captivity, and that tend to be cheaper, have a higher probability of being introduced into the wild (Robinson, 2001; Cassey et al., 2004b;Blackburn, Lockwood & Cassey, 2009). In general, among bird species, Psittaciformes have a high probability of transport and introduction outside their native range (Lockwood, 1999; Lockwood, Brooks & McKinney, 2000; Blackburn & Duncan, 2001; Duncan, Blackburn & Cassey, 2006), being traded as much as 14 times more often than other avian orders (Bush, Baker & MacDonald, 2014). Moreover, parrots in general have a wide diet breath, and research suggests that diet breath and migratory tendencies can explain the success of established exotic populations of parrots (Cassey et al., 2004a). Consequently, 10–16% of all parrot species have established exotic populations around the world (Cassey et al., 2004b; Menchetti & Mori, 2014).

As with any introduced invasive species, there are concerns over the possible negative impacts that exotic parrots may have on the economy and the native ecosystems (Pérez-Rivera & Vélez-Miranda, 1980), which has been recently reviewed by Menchetti & Mori (2014). These include damage to crops, damage to the electrical infrastructure, transmission of diseases, competition, and hybridization. Damage to crops is perhaps the biggest negative impact that parrots cause in both their native and introduced range, with substantial economic losses (e.g., González, 2003). In addition, in Florida (USA), monk parakeets have been reported to cause electric shortages while building nests in electric towers, costing as much as USD 585,000 for repairs (Avery et al., 2006). To our knowledge, the only invasive species that has caused electrical shortages in Puerto Rico has been the invasive green iguana (*Iguana iguana*; Falcón et al., 2013), and we have not observed nests of monk parakeets in electrical towers. Moreover, wild and captive Psittaciformes are known vectors of avian and diseases and parasites, and also human diseases (Clark, Hume & Hayes, 1988; Orosz et al., 1992; Magnino et al., 1996; Mase et al., 2001; Azevedo, 2014; Done & Tamura, 2014; Briceno et al., 2017). In terms of competition, antagonistic interactions have been reported in parks and urbanized areas where introduced parrots aggressively displace native birds at garden feeders, they can consume unripe fruits before other native birds have access to them (because native species only eat them when they are ripe; Snyder, Wieley & Kepler, 2007; Lin Neo, 2012; Le Louarn et al., 2016). Competition for cavities, on the other hand, may be a cause of concern, because there are native species which depend on this scarce resource, and psittacine secondary cavity nesters can be aggressive competitor (Snyder, Wieley & Kepler, 2007; Strubbe & Matthysen, 2009a; Orchan et al., 2012; Mori et al., 2017). However, populations of cavity nesters (e.g., *Amazona spp* and rose-ringed parakeets) are relatively low, and the most successful psittacine species in Puerto Rico build their own nests. Finally, based on the distribution of *Amazona spp.* in Puerto Rico, hybridization with the endemic Puerto Rican amazon is unlikely at the moment, but it is a latent possibility as both endemic and introduced populations continue to grow and expand their range. For example, *Amazona oratrix* and *A. aestiva* are known to hybridize in the wild when they co-occur (in their introduced range; Martens, Hoppe & Woog, 2013).

Another aspect to consider is that the three species of endemic parrots in Puerto Rico were once much more common and used to occupy many of the areas and habitats now occupied by exotic species. In fact, the extinct Puerto Rican parakeet (*Psittacara maugei*) was so abundant and widespread that it was, in part, hunted down to extinction because of the damages it caused to the agricultural sector (Olson, 2015), as was the Carolina parakeet (Saikku, 1990). Thus, the dramatic population decrease and range reduction of the Puerto Rican parrot, as well as the extinction of the other endemic parrot and parakeet left empty niches. Perhaps one of the most important functions performed by the endemic parrots were seed predation and seed dispersal. Parrots are known to incur in such functions (Norconk, Grafton & Conklin Brittain, 1998; Francisco, Lunardi & Galetti, 2002; Blanco et al., 2015; 2016; Blanco, Hiraldo & Tella, 2017), and both seed predation and seed dispersal have important implications for ecosystems dynamics worldwide, and help regulate plant recruitment, competition, and population structure (Howe & Smallwood, 1982; Hamrick, Murawski & Nason, 1993; Nathan & Muller-Landau, 2000). Unlike other birds, parrots have specialized bills that allows them to access resources, such as hard seeds, that are often not available to other animals, and they often predate on them in the wild. For example, the white-winged parakeet is a seed predator of the Panama tree (*Sterculia apetala*), an introduced species in Puerto Rico. But they also predate seeds of the pink trumpet tree (*Tabebuia heterophylla*), an abundant native species on the island. Likewise, blue-and-yellow macaws predate on the seeds of *Swietenia spp.*, which are exotic trees whose seeds are too large for other species of birds. On the other hand, as generalist frugivores, parrots also act as seed dispersers via endozoochory (Blanco et al., 2016). For example, white-winged parakeets are known to disperse *Ficus spp.*, which are small- and hard-seeded plants (WF, pers. obs.). Research should focus on the ecological role of introduced psittacine species in order to assess the impacts, neutral, positive and/or negative, that they are having in their introduced range.

Currently, no species of Psittaciformes is considered illegal in Puerto Rico by the Department of Natural and Environmental Resources (Departamento de Recursos Naturales y Ambientales de Puerto Rico, 2003). Trapping of exotic birds established in Puerto Rico is allowed by the DNER for exportation only, and the sale in local markets is prohibited. A permit is required for trapping and exporting birds, and a report must be filed at the end of the activity. We had access to such reports from the DNER, and the most commonly trapped bird was the white-winged parakeet. However, rarely the number of individuals trapped was reported. Moreover, information about the exportation process is not provided. We recommend that the DNER adjusts the protocol and collects detailed information about the locality of trapping, quantity and species of birds trapped, and the final destination of the birds. Such information may provide insights into how harvesting affects the population growth of these birds and therefore have management implications.

Although selling some of the wild trapped species of parrots in the local pet market is illegal, it is common to find people selling them, especially on the internet. Particularly common are budgerigars, lovebirds, white-winged parakeets and monk parakeets. The rose-ringed parakeet is gaining popularity and it is relatively easy to acquire them. Therefore, we recommend that management agencies should prohibit or limit the sale and possession of psittacine species that are prone to establish outside their native range, and focus on the trade of Psittaciformes (and other species) in order to prevent more escapes and thus control propagule pressure and range expansion via introductions to new areas.

If control and management is necessary and/or desirable, there are several methods available. Public education about the potential effects of releasing exotic animals, and breeding controls are important for preventing releases into the wild. When direct management is necessary, there are three options: 1) trapping and exporting birds, 2) birth control chemosterilants such as Diazacon^™^ (Yoder et al., 2007; Avery, Yoder & Tillman, 2008; Lambert et al., 2010), and 3) culling (lethal). However, control efforts, especially lethal ones, may be hindered by members of the public, who usually protest these actions as parrots have a high aesthetic value (Avery & Tillman, 2005).

### Ecology of Brotogeris versicolurus in Puerto Rico

Diet observation of the white-winged parakeet coincide with previous reports on the island (Pérez-Rivera et al., 1985), are consistent to those reported elsewhere in their invasive and native range (Brightsmith et al., 2017), and show that although they use native species, the majority of the plant species consumed are exotic ones. Nesting phenology observations in the Metropolitan Area of San Juan are consistent with the findings of the population in San Germán (Tossas, Colón & Sanders, 2012), and in Florida (Brightsmith et al., 2017). Moreover, we found an unidentified species of mite (probably *Pararalichus*), which has been reported on *B. versocolurus* in its native range, along with *Aralichus cribiformes* and *Echinofemur sp.* and *Rhytidelasma sp.* (Atyeo, 1989). In most cases, transfer of mites occur by physical contact between conspecifics, but there are cases of inter-species transfer (Dabert & Mironov, 1999; Hernandes, Valim & Pedroso, 2016). However, these, as most feather mites, are considered ecto- commensals and feed on the oils produced by the birds (Blanco et al., 2001).

Calculating the population growth rate based on the population size estimate of Tossas, Colón & Sanders (2012) and our first population estimate yields results of positive growth rate consistent with those found during our study period. Comparing the historic number of parakeets in the study populations with data from this study, and their population growth rates, suggest that the population exhibited a lag-phase, and was growing exponentially by 2010.

The resulting population decrease in 2010 was probably due to emigration and the establishment of a new roosting site in the Río Piedras Botanical Garden, where parakeets would forage, but did not roosted before (5.6 km from the San Patricio roosting site). Around 800 parakeets were seen roosting at this new location. If we assume that the parakeets at the new roosting site originated from the San Patricio population, then a minimum of 200 fledglings were produced (considerably lower than the production in San Germán), and about 300 pairs abandoned the San Patricio roost. Another roosting site was established in 2012 at the University of Puerto Rico, 5.5 km from San Patricio, and 1.5 km from the Botanical Garden population.

Population growth rates of species of birds regarded as invasive are usually exponential, and often exhibit a lag-phase (Sakai et al., 2001; Blackburn, Lockwood & Cassey, 2009) and we have observed both of these patterns in the Puerto Rican populations of *B. versicolurus* (and *M. monachus).* Exotic species that exhibit high population growth rates may be more likely to establish viable populations (Ehrlich, 1986; Moulton & Pimm, 1986; Pimm, Jones & Diamond, 1988; Pimm, 1989; Cassey, 2001). Additionally, as the size of species population increases, so does the geographic distribution, so we can expect that population growth will probably result in spread (although this is not always the case; Gaston et al., 2003; Gaston, 2003; Blackburn, Cassey & Gaston, 2006). Indeed, in the yellow-headed blackbird (*Xanthocephalus xanthocephalus*), a native species in the USA, the eastward dispersal of the species was linked with the population growth rate (Veit, 1997). It has been proposed that populations of birds exhibiting a positive population growth rate have a proportional increase of long dispersers (vagrants) to the increase in population size and thus, incrementing the chances of the species invading new ranges (Veit, 1997; 2000). Our data and observations suggest that the rapid increase and high number of parakeets in the San Patricio population led to the emigration and establishment of new roosts, therefore expanding their range. This is supported by the lag between population increase at the largest roosting sites on the island, and the island-wide population increase.

Migration to and from the roosting sites is a factor that can affect our population growth estimates, and bias our results since we are not considering migration and we assume that the changes in populations are due to birth and death rates. Our surveys in the vicinity of both study populations suggested that they were they only existing roosting sites in the areas, a fact that was confirmed by Tossas, Colón & Sanders (2012) for the San Germán population. During our surveys, we were able to detect the establishment of a new roosting site, which probably originated from the San Patricio population, so emigration is a factor regulating population size. Although white-winged parakeets can exhibit yearly variations in terms of the number of individuals at the roost site, the variation is low within the peak of the breeding and non-breeding seasons (Tossas, Colón & Sanders, 2012), as confirmed by our weekly estimates. Given this, and that we were able to detect the establishment of the new roosting site, which likely explains the population decline in San Patricio, we are confident that our methods and estimates are robust.

Various factors may have contributed to the success of white-winged parakeets in Puerto Rico. Firstly, propagule pressure may have been an important factor leading to the introduction and establishment during the 1960’s and 1980’s, when they were popular and cheap. According to the CITES Trade Database, the main exporter to the USA (and therefore Puerto Rico) was Peru, with 2,832 white-winged parakeets exported in a single shipment in 1985. Moreover, because they nest in termite mounds, which are highly abundant in the secondary forests of Puerto Rico and parks, white-winged parakeets are not limited by the scarcity of tree cavities as in other parrot species. In addition, no predation observation have been made, indicating that predation pressure is low, and the potential predator pool is rather small (including *Falco peregrinus, F. columbarius, F. sparverius, Acipiter striatus* and *A. cooperii*; Brightsmith et al., 2017). Furthermore, trapping seems to supply the local (informal) pet market, rather than serving for exportation, further exacerbating releases into the wild. Finally, the white-winged parakeet seems to have a wide diet breath.

### Conclusions

Our data shows that most Psittaciformes introduced through the pet trade to Puerto Rico, and that established populations, are still present and persist in the wild. Based on population sizes and range, white-winged parakeets and monk parakeets are indeed the most successful Psittaciformes in Puerto Rico. The data suggest that white-winged parakeets are very prolific, are aggressively expanding their range, especially in coastal areas, and we expect them to keep growing exponentially and expanding into new, unoccupied areas. The same is probably true for monk parakeets. A species of special concern is the rose-ringed parakeet (*Psittacula krameri).* This species is highly invasive elsewhere, and has caused negative ecological impacts (Butler, 2003; Strubbe & Matthysen, 2009a,b; Kumschick & Nentwig, 2010; Newson et al., 2011; Sa et al., 2014; Le Louarn et al., 2016). Therefore, special attention should be given to this species, and others that have invasiveness potential, with the aims of preventing the sale and possession and ultimately prevent releases to the wild and subsequent establishment.

Although unlikely at the moment, as populations of the endemic and endangered Puerto Rican parrot increase in size and distribution, it is likely that they will interact with exotic species. Thus, special consideration and management priority should be given to species with populations established near wild populations of Puerto Rican parrots, especially other *Amazona spp.*, to prevent competition for food and nesting resources, hybridization and transmission of diseases.

It is worth mentioning that in late 2017 Hurricane María, a category four hurricane, hit Puerto Rico with a south-to-north trajectory, and undoubtedly caused negative impacts on the parrot populations in the island. For example, dozens of dead white-winged parakeets were observed around a recent roost located in Río Piedras (municipality of San Juan; Fig. 1f), and at least one blue-and-yellow macaw was found dead in Guaynabo. Despite this, numerous flocks of parakeets and at least eight macaws were seen after the hurricane, so a proportion of individuals of these species survived. It is highly possible that other parrots experienced mortality related to the hurricane. Indeed, the endemic Puerto Rican parrot suffered very high hurricane-related losses in the east of the island (R Valentín, pers. comm.). Another negative effect as a result of the hurricane is the lack of food resources due to the massive exfoliation of food plants. For example, blue-and-yellow macaws, which usually forage high on trees and palms, were observed eating flowers on shrubs as low as 1.5 m from the ground due to the lack of food. A follow-up study on the status of Psittaciformes after the hurricane is recommended, as it is possible that the negative effects may result in the extirpation of some of the species, especially those with small population sizes.

Finally, many of the species of Psittaciformes found in the wild in Puerto Rico are vulnerable or endangered in their native range, and introduced populations provide the opportunity to conduct experiments and/or to explore management techniques that otherwise would be impossible to perform in their native habitat, both aspects which may aid in the conservation of Psittaciformes in their native habitat around the world.

## Acknowledgements

We want to especially thank Rafael D. Rodríguez, Linda Ortíz and Rebecca Hernández for their help in the field during the white-winged parakeet roost counts. We also thank Marilyn Colón of the DNER, for her help during permit application process, and for providing unpublished reports on exotic bird trapping in Puerto Rico, and Ricardo Valentín of the DNER for sharing information about the state of the Puerto Rican Parrot after Hurricane Maria. We also thank and Noramil Herrera, and Alberto Mercado (DNER) for helping in the search of literature resources. Julio A. Salgado of the Puerto Rico Ornithological Society provided assistance in sharing the online form for reporting Psittaciformes in Puerto Rico among different local groups, and also provided information on the current status of psittacine birds on the island, and photos, for which we are extremely grateful. We also thank Yoly Pereira, Pedro Santana, Sonia Longoria, and Dinath Figueroa for allowing us to use their photos. The PR-Louis Stokes Alliance for Minority Participation Bridge to the Doctorate Fellowship (HRD-0601843), the Ronald E. McNair Program of the University of Puerto Rico at Humacao (P217A070213), and the Center for Applied Tropical Ecology and Conservation (HRD-0734826) provided funding to WF in partial support of this project.

